# Mucin hydrogel glyco-modulation to investigate immune activity of mucin glycans

**DOI:** 10.1101/2019.12.18.880757

**Authors:** Hongji Yan, Morgan Hjorth, Benjamin Winkeljann, Illia Dobryden, Oliver Lieleg, Thomas Crouzier

**Affiliations:** Division of Glycoscience, Department of Chemistry, School of Engineering Sciences in Chemistry, Biotechnology and Health, KTH, Royal Institute of Technology, AlbaNova University Center, 106 91 Stockholm, Sweden; Department of Mechanical Engineering and Munich School of Bioengineering, Technical University of Munich, Boltzmannstrasse 11, 85748 Garching, Germany; Division of Surface and Corrosion Science, Department of Chemistry, School of Engineering Sciences in Chemistry, Biotechnology and Health, KTH Royal Institute of Technology, Drottning Kristinas väg 51, 10044 Stockholm, Sweden

## Abstract

Mucins are multifunctional glycosylated proteins that are increasingly investigated as building blocks of novel biomaterials. Once assembled into hydrogels (Muc gels), mucins were shown to modulate the recruitment and activation of immune cells and avoid fibrous encapsulation *in vivo*. However, nothing is known about the early immune response to Muc gels. This study characterizes the response of macrophages, important orchestrators of the material-mediated immune response, over the first 7 days in contact with Muc gels. The role of mucin-bound sialic acid sugar residues was investigated by first enzymatically cleaving the sugar, then assembling the mucin variants into covalently crosslinked hydrogels with rheological and surface nanomechanical properties similar to non-modified Muc gels. Results with THP1 and human primary peripheral blood monocytes-derived macrophages were strikingly consistent and showed that Muc gels transiently activate the expression of both pro-inflammatory and anti-inflammatory cytokines and cell surface markers, with a maximum on the first day and loss of the effect after 7 days. The activation was sialic acid-dependent for a majority of the markers followed. The pattern of gene expression, protein expression, and functional measurements did not strictly correspond to M1 or M2 macrophage phenotypes. This study highlights the complex early events in macrophage activation in contact with mucin materials and the importance of sialic acid residues in such a response. The enzymatic glyco-modulation of Muc gels appears as a useful tool to help understand the biological functions of specific glycans on mucins which can further inform on their use in various biomedical applications.

## Introduction

Breakthroughs in materials engineering have accelerated the use of biomaterials in both preclinical and clinical applications, including engineered cell microenvironments^1–3^, drug delivery^4^, tissue engineering^5^, and immunoengineering^6^. A new class of biomaterials has emerged that is not designed to be ‘biologically inert’ but rather to deliver a provision of cues to surrounding cells resulting in improved material performance^7,8^. This approach is particularly valuable when considering that the immune response to implanted biomaterials can help suppress or modulate the immune cascades to avoid acute inflammation or subacute inflammation. These materials find applications in regenerative medicine where hyperactivity of immune cells in a damaged tissue is suppressed by the material to promote the healing process^9,10^, or in cancer therapy^11–13^ and novel vaccine therapies^14^, where a complex immune-modulation from the material can help eradicate diseased cells and promote healthy cells via a myriad of coordinated intra- and extra-cellular signaling pathways.

Mucin glycoproteins are emerging as attractive building blocks to assemble such bioactive materials^15,16^, driven by advances in our understanding of their structure and biological functions. Mucins are a family of glycosylated proteins, and more than 50% of their mass is composed of *O*-glycans. Mucins are found bound to the cell membrane as part of the glycocalyx^17^, or secreted to form the mucus gel protecting the epithelium against irritants, pathogens, and to provide hydration and lubrication^15,18^. Mucins undergo dynamic changes in physiological conditions to maintain homeostasis, as well as in the initiation and development of diseases. In many types of tumors, transmembrane mucins are altered in their expression level^19^ and glycosylation pattern marked by a general reduction of glycosylation^20^ and truncation of *O*-linked glycans^21^. In mucinous carcinomas, secreted mucins surround the tumors protecting them from cancer drugs and immune cell infiltration both physically and biochemically^22^. Mucins are immunologically active through the binding of their sugar residues to lectin-like proteins on the surface of immune cells^18,23,24^. The muc2 mucins found in the gut can imprint dendritic cells tolerance^23^ and in contrary, can activate non-stimulated dendritic cells in a concentration-dependent manner^25^.

We have recently shown that covalently crosslinked mucin hydrogels (Muc gels) made of bovine submaxillary mucins (Muc) modulate the foreign body response *in vivo*. Those hydrogels caused a broad dampening effect of cytokine expression in macrophages harvested from the explanted gels and their corresponding peritoneal cavity, and the absence of fibrous encapsulation after 21 days^26^. This discovery suggests that biomaterials containing mucins or mucin-like molecules could be used as implantable hydrogels and coatings that can evade fibrosis and ensure the long-term function of the devices. However, fibrous capsule formation is typically initiated after ∼ 2 weeks of implantation^26^, and the earlier events that led to such effects are unknown. In addition, none of the features of mucins essential for their immune-modulating properties were clearly identified.

We address these limitations herein by investigating the response of undifferentiated macrophages (M0) derived from monocyte cell line THP1 and human primary peripheral blood monocytes when cultured on the surface of Muc and tMuc gels over 7 days. The glyco-modulated Muc gels (tMuc gels) are used to highlight the role of mucin-bound sialic acid sugar residues in the immune-modulating effect. We focus the study on macrophages since material-immune interactions are predominantly orchestrated by macrophages *in vivo*, owing to their heterogeneity and plasticity^8^. Unlike other terminally differentiated cells, macrophages can sense cues from their environment and undergo dynamic changes^27^, either to fight against pathogens^8^, or to contribute to tissue healing via directing stromal cell recruitment and differentiation to maintain tissue homeostasis^27^. And in tumors, tumor-associated macrophages (TAMs) contribute to a tumor immunosuppressive microenvironment that protects tumor cells^28^. The cues directing macrophage phenotype and function can originate various microenvironments, including from mucus gels that were shown to determine macrophage phenotype and functions in the lung^29^, and are hypothesized to do so in the gut and TAMs of mucinous cancers. A detailed study of how materials and, in particular, mucin-materials can modulate the activity of macrophages could help us understand the many processes macrophages are involved in.

## Results

### The glycan composition of Muc gels can be modulated by enzymatic treatment without compromising the mechanical properties of the gels

To prepare mucin hydrogels (Muc gels), we introduced tetrazine (Tz) and norbornene (Nb) functionalities to bovine submaxillary mucin (Muc) molecules as previously described^26^. After being mixed in solution, the Muc-Tz and Muc-Nb formed a covalently crosslinked hydrogel through an inverse electron demand Diels-Alder cycloaddition reaction (**Figure 1A, B**). To investigate the role of sialic acid in the response of macrophages, we cleaved sialic acid residues by treating Muc-Tz (tMuc-Tz) and Muc-Nb (tMuc-Nb) with neuraminidase (**Figure 1B**). We show by anion exchange chromatography that about 60% of all sialic acid residues were removed after neuraminidase treatment (**Figure 1C, D**). This incomplete removal of sialic acid could be due to the inaccessibility a fraction of the sialic acid residues or to the specificity of the neuraminidase used. However, given that we obtained an even removal efficiency for the modified and unmodified mucins, the presence of Tz and Nb, which we hypothesise to be located on the mucin protein backbone (**Figure SI1**), does not seem to be responsible for this incomplete sialic acid removal.

**Figure 1.**
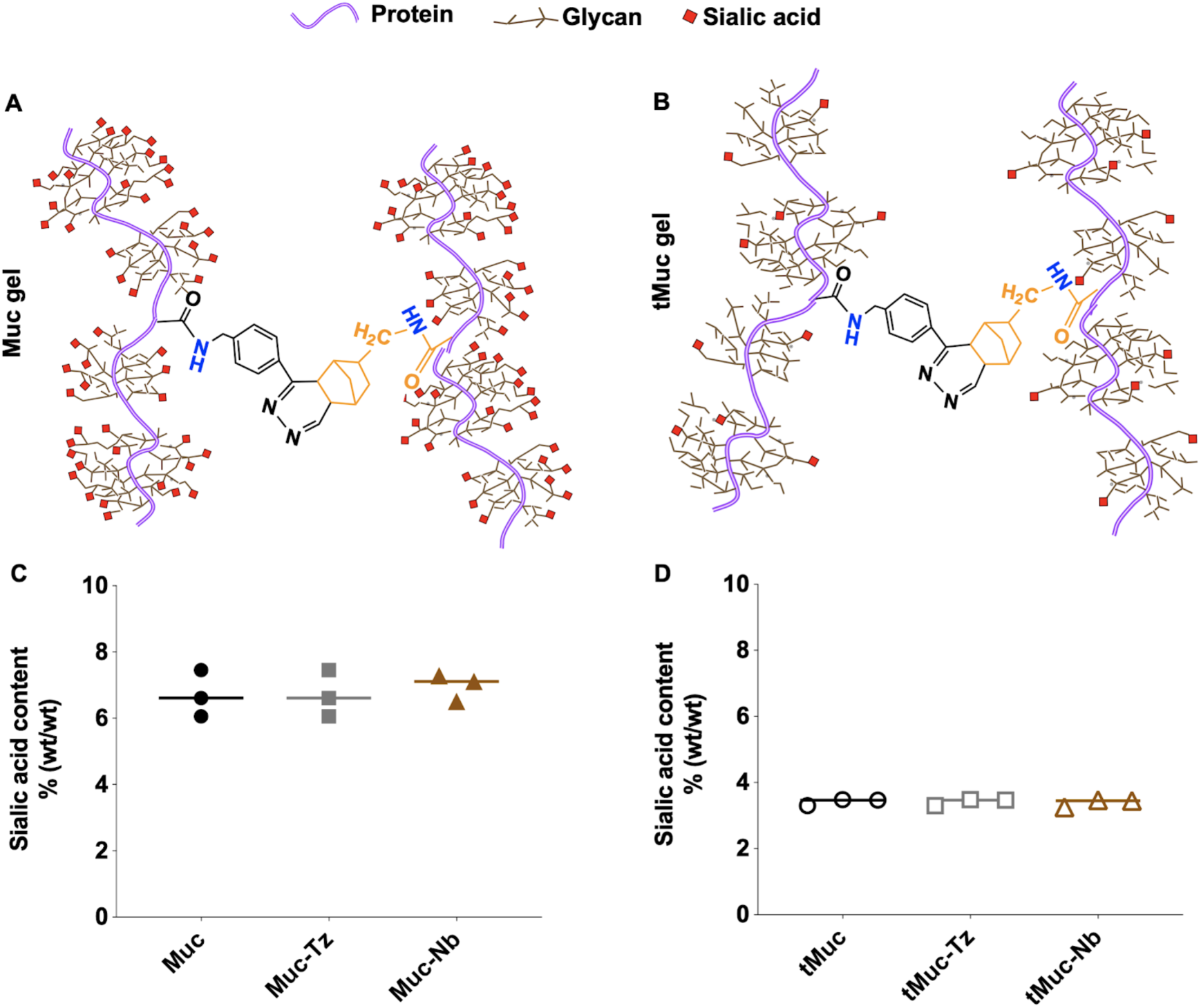
Muc gels crosslinking reaction and mucin glycan modification. Representation of the crosslinking reaction of Muc gels (A) and tMuc gels (B, neuraminidase treated). Quantification of sialic acid residues on Muc, Muc-Tz, and Muc-Nb (C) and neuraminidase-treated tMuc, tMuc-Tz, and tMuc-Nb. The data points are obtained from measurements of *n* = 3 independent samples.

We then tested whether the enzymatic treatment would compromise the rheological properties of the hydrogels; such an effect could influence the macrophage response to the material^30^, and make the contributions of sialic acid difficult to infer. Muc-Tz and Muc-Nb solubilized in PBS were mixed and then subjected to oscillatory rheology measurements over time. Both the loss (*G*’’) and storage (*G*’) moduli rapidly increased, and initially the response was dominated by *G*’’ (indicating the presence of a viscoelastic solution) (**Figure 2A**). However, ∼5 min after mixing, the response became dominated by *G*’ indicating the presence of a viscoelastic solid thus confirming successful gel formation (**Figure 2A, insert**). Both, storage and loss modulus reached a plateau-like state after ∼60 min. This plateau value of G’ was ∼ 10 kPa, which is several orders of magnitude higher than the elastic modulus of an uncrosslinked, entangled mucin solution^31^. A frequency sweep performed after the viscoelastic moduli have reached plateaus demonstrated that the system appears indeed to be efficiently and covalently cross-linked (**Figure 2B**). Gels with similar viscoelastic properties were obtained when using a complete cell culture medium to dissolve the mucins, suggesting the medium did not interfere with the crosslinking reactions occurring between Tz and Nb (**Figure SI2**). Importantly, neuraminidase-treated tMuc-Tz and tMuc-Nb also reacted to form hydrogels and showed a rheological behavior and calculated mesh size^5^ (ξ) similar to those of untreated Muc gels (**Figure 2C, insert, and D; Table 1**).

**Table 1.**
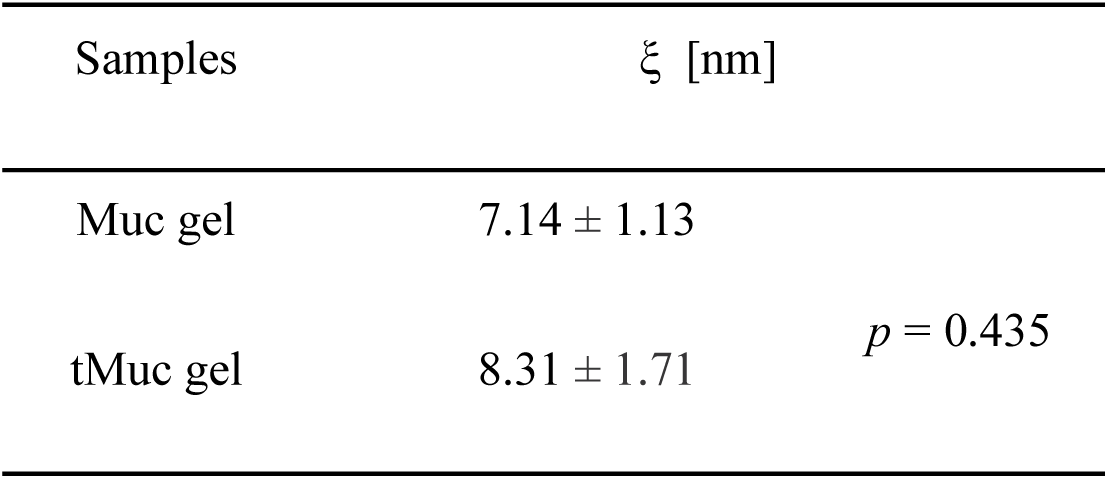
Mesh size values estimated from the rheology data shown in **Figure 2** (n=3)

**Figure 2.**
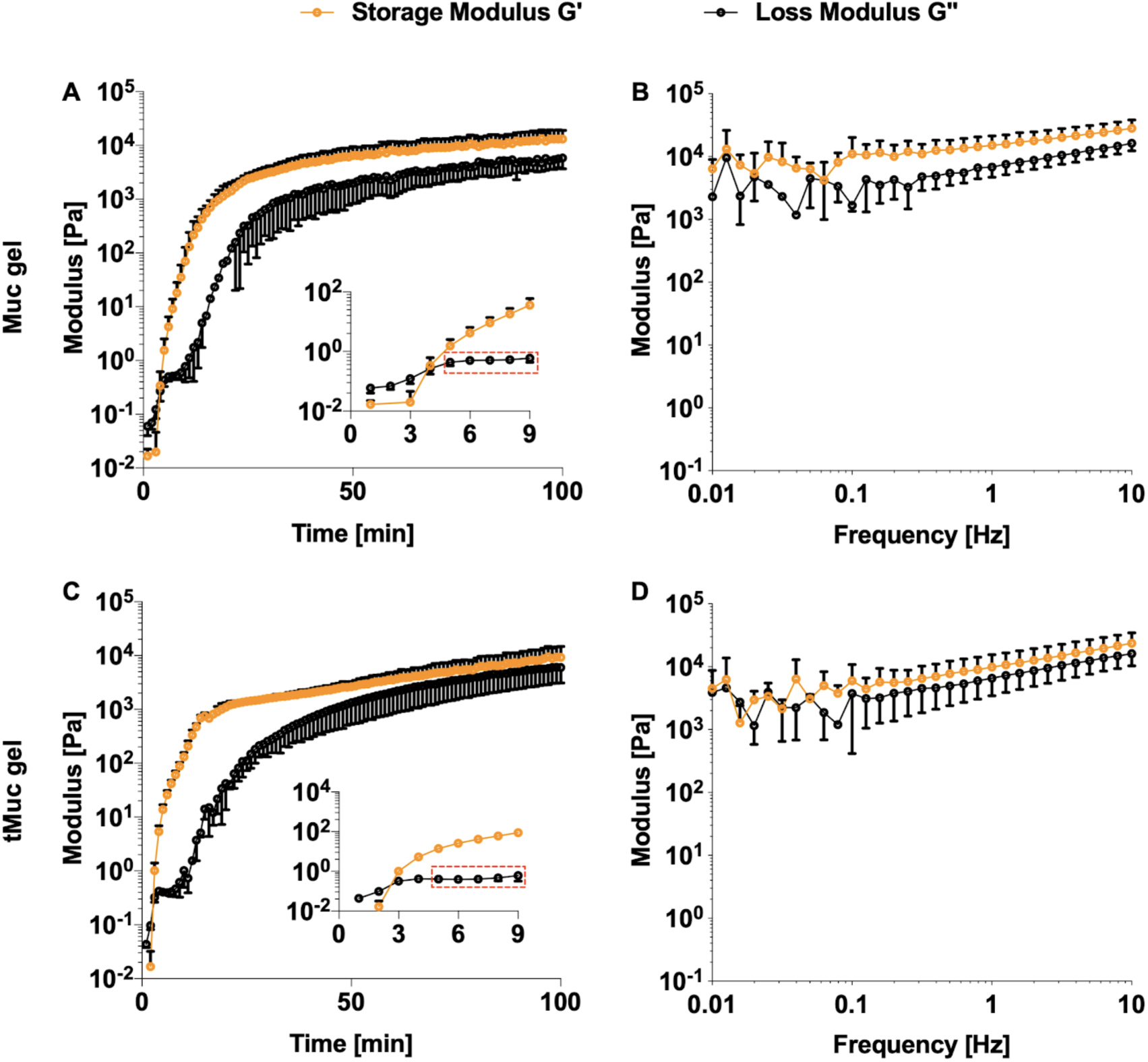
Rheological characterization of Muc gels and tMuc gels. Time-dependent rheological measurements of the mixed Muc-Tz and Muc-Nb (A) or tMuc-Tz and tMuc-Nb (C) in PBS; Final frequency-dependent viscoelastic moduli of the crosslinked Muc-gel (B) and tMuc gels (D). The error bars denote the standard deviation as obtained from measurements of *n* = 3 independent samples.

We further characterized the nanomechanical surface properties of hydrated Muc gels and tMuc gels by atomic force microscopy (AFM)-based nanomechanical surface mapping with the tip submerged in PBS. The measurement is complementary to the bulk rheometer measurements and allows us to reveal the surface heterogeneity in nanomechanics in a range of the AFM tip’s radius^32^. We recorded force volume maps for both approach (a combined elastic and viscous contribution) and retraction regimes (mainly elastic contributions). The average elastic modulus (**Figure 3A, B**) calculated from the elastic modulus maps (Fi**gure SI3, 4**) showed no difference between Muc gels and tMuc gels. There was also no difference in the stiffness (**Figure 3C, D**) calculated from the slopes in the repulsive part of the force curves, which are independent of contact models^33^.

**Figure 3.**
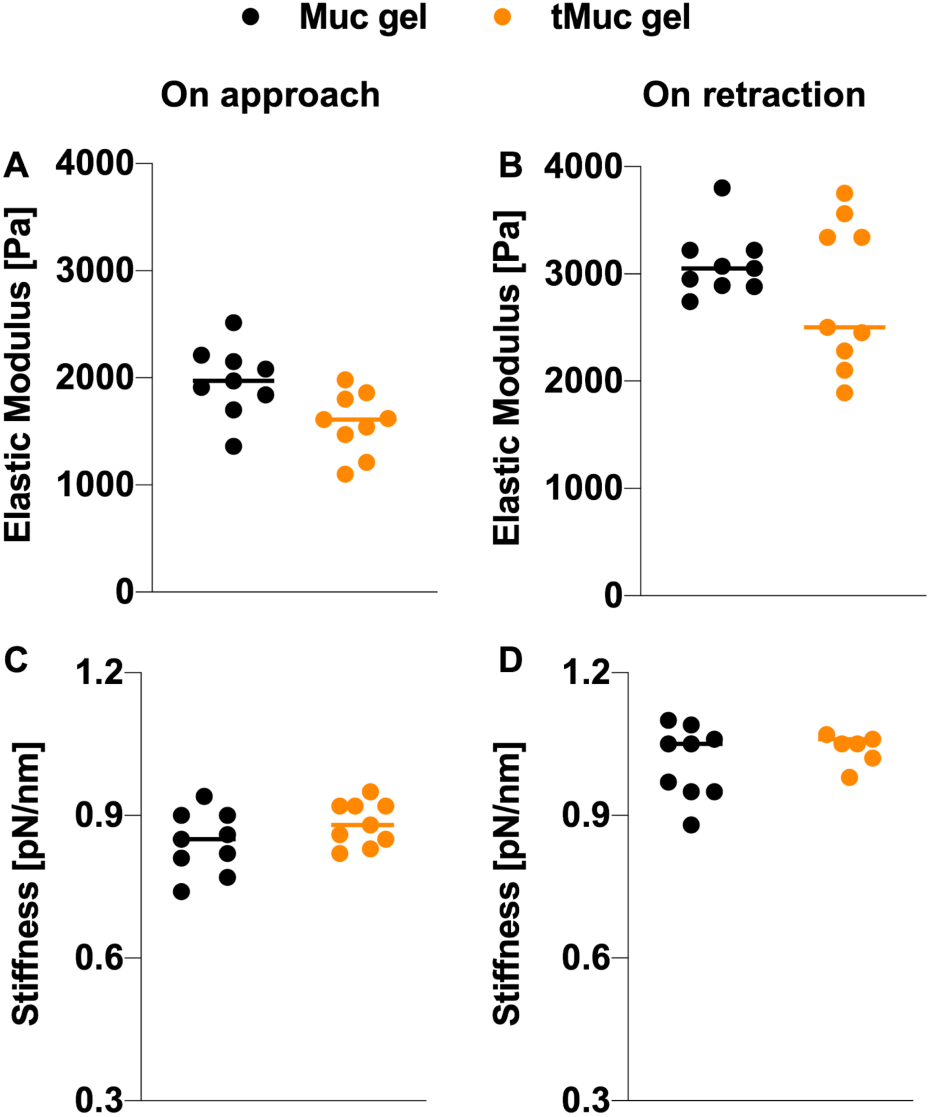
AFM nanomechanical characterization of Muc gels and tMuc gels. Elastic moduli (A, B) and stiffness (C, D) of Muc gels and tMuc gels were obtained by AFM-based force volume mapping for both approach (elastic, viscous and viscoelastic contributions) and retraction regimes (mainly elastic contribution). n=9.

### THP-1 derived macrophages type 0 are activated by Muc gels in a sialic acid dependent manner

To investigate the early response of macrophages to mucin materials we first used macrophages type 0 differentiated from human monocyte cell line THP1 (THP1-M0) by incubation with phorbol 12-myristate 13-acetate (PMA, 150 nM) for 3 days followed by incubation in a complete cell culture medium without PMA for 1 day. After differentiation, the cells became adherent on tissue culture polystyrene (TCP) and expressed increased levels of CD36 and CD71 macrophage markers^34^ compared to THP1 monocytes (**Figure SI5**). We seeded THP1-M0 on tissue culture polystyrene (TCP), Muc gel and tMuc gels and cultured them over a period of 7 days. The THP1-M0 did not adhere strongly, did not spread, and formed clusters within hours on both Muc gels and tMuc gels (**Figure 4**).

**Figure 4.**
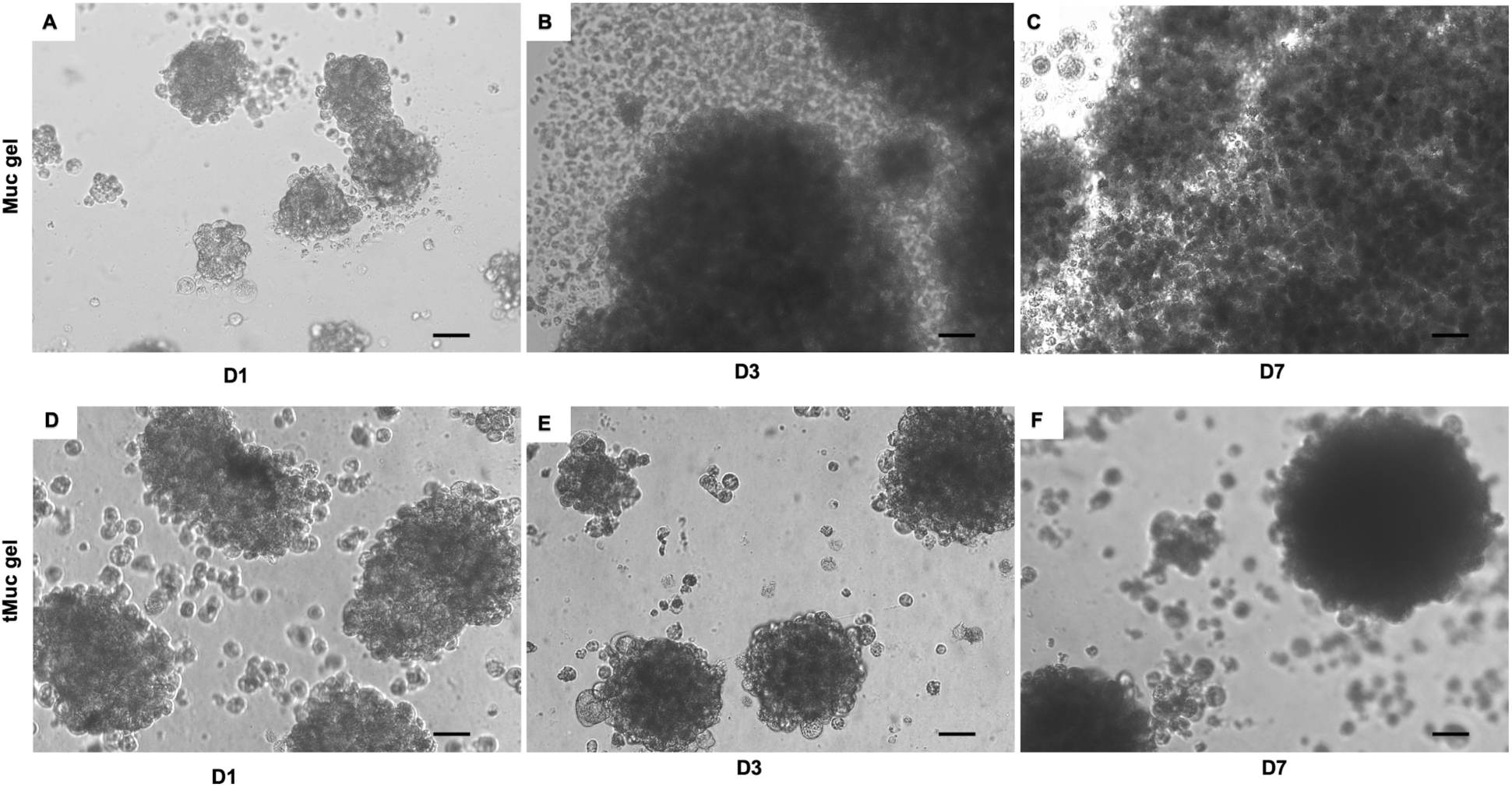
Representatives of phase contrast images of THP1-M0 cultured on Muc gel and tMuc gel on day 1, 3, and 7 (D1, D3, and D7). Scale bar = 50 µm.

We then ask whether undifferentiated M0 macrophages would be activated and be polarized when in contact with Muc gels. Historically, macrophages have been broadly classified into proinflammatory phenotype (M1) that is stimulated by pro-inflammatory signals, such as interferon-*γ* (IFN-*γ*) or microbial products lipopolysaccharide (LPS)^35^, and alternatively activated (M2) that is stimulated by signals from basophils, mast cells, and other granulocytes^35^. M1 cells have higher capacity in antigen-presenting, and enhancing Th1 differentiation of lymphocytes that produces the proinflammatory signals^35,36^. M1 cells also harm adjacent cells via producing toxic reactive oxygen species (ROS) and escalating the proinflammatory responses^37^. M2 also constantly express scavenger and mannose receptors and release anti-inflammatory cytokines, i.e., IL-10^35^.

We measured the gene expression of 11 pro- and anti-inflammatory macrophages markers by RT-PCR (**Table S1)**. There was no significant difference in expression of the majority of markers over 7 days between non-adhesive and adhesive TCP (**Figure SI6**) but with a slight activation of THP1-M0 for some cytokines (*i.e., CXCL10, CXCL8*, and *CCL2*) on adhesive TCP. We thus selected adherent TCP as reference material even though M0 macrophages adhere to TCP and not Muc gels. Both pro-inflammatory *CXCL10, CXCL8, TNFa, CCL2IL1B, VEGFA*, and anti-inflammatory *IL1RN* cytokines were upregulated on the first day, then followed by a decrease on day 3 and 7 in THP1-M0 cultured on Muc gels when compared to TCP and tMuc gels (**Figure 5**). *IL-10*, an anti-inflammatory cytokine, showed a unique gene expression pattern with a later activation on day 3, followed by a decrease on day 7. M2 macrophages surface markers (*Tgm2, MCR1*) did not change compared to TCP control but was downregulated on day 7. M1 surface markers *CD64* was significantly downregulated in THP1-M0 cultured on Muc gels. The partial removal of sialic acid had a profound effect on the response of THP1-M0 macrophages when compared to Muc gels. For nearly all markers, THP1-M0 cultured on tMuc gels led to little or no activation on day 1 and 3, in contrast with the strong transient activation observed in THP1-M0 cultured on Muc gels. Exceptions were for CD64, which was significantly higher for macrophages cultured on tMuc gel compared to Muc gel on day 1 and MCR1, which was not affected.

**Figure 5.**
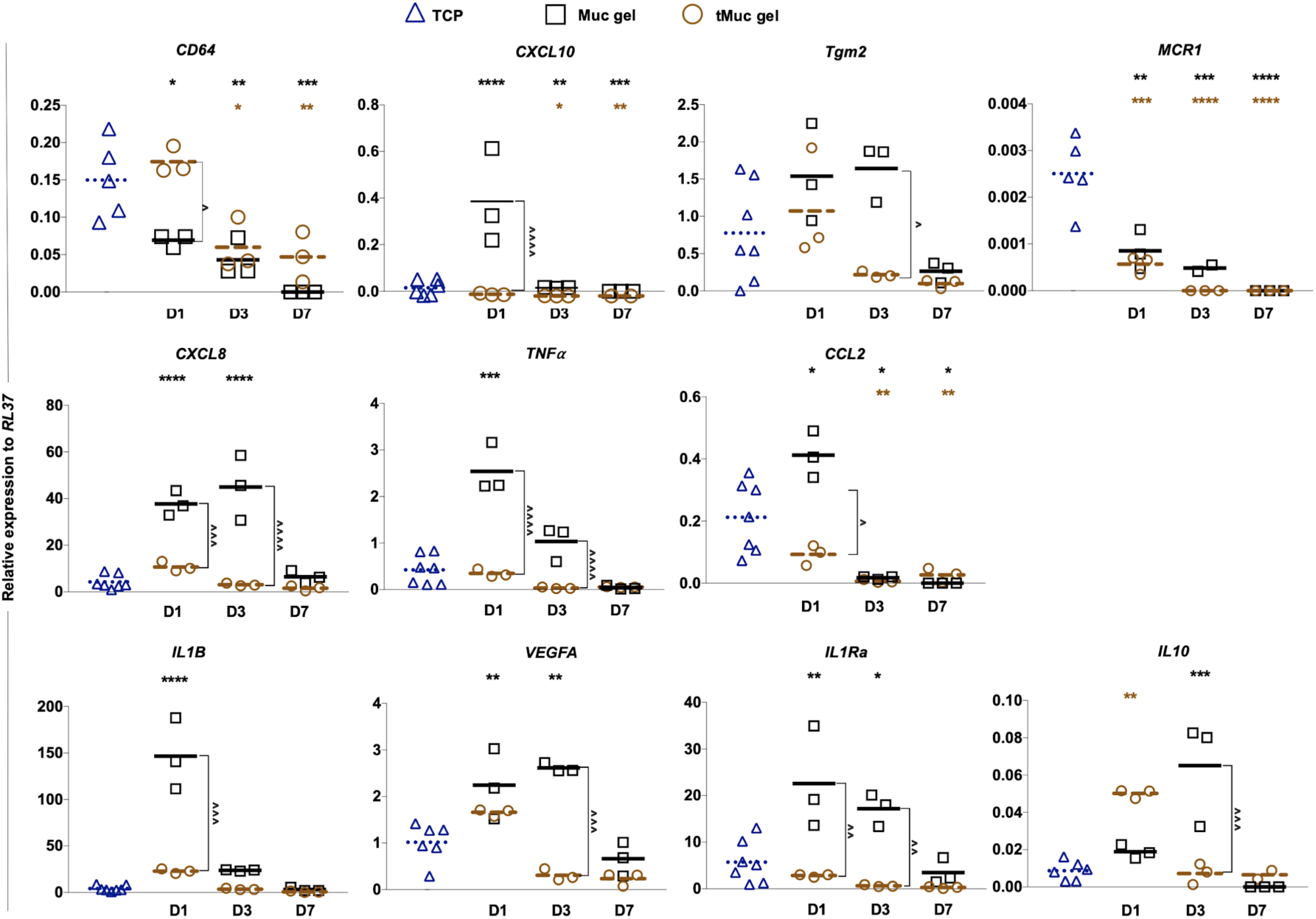
Gene expression in THP-1 derived macrophages type 0 (THP1-M0) after being cultured on tissue culture polystyrene (TCP), Muc gels and Sialidase treated Muc gels (tMuc gel) on Day (D) 1, 3 and 7. The data points denote the mean of relative gene expression to *RPL-37* obtained from three independent experiments with duplicates. Statistical significance was calculated by One-way ANOVA test by Prism 8.0. Black*, Brown*, and Black^ indicates the comparison between Muc gels vs TCP, tMuc gels vs TCP, and Muc gel vs tMuc gel individually. ‘*’, ‘**’, ‘***’, and ‘****’ indicate p values of < 0.05, 0.01, 0.0005, and 0.0001, respectively.

We confirmed the gene expression by measuring the expression of four intracellular cytokines at the protein level by FACS. As shown in **Figure 6**, the results were consistent with gene expression with IL1Ra, IL-1B, CXCL8 in THP1-M0, all significantly upregulated in THP1-M0 cultured on Muc gels when compared to TCP. Reduction in sialic acid content also led to an inhibition of the transient activation of the macrophages. Of note, *IL10* protein expression was decreased in macrophages cultured on tMuc gel compared to Muc gel which is in discrepancy with the gene expression data.

**Figure 6.**
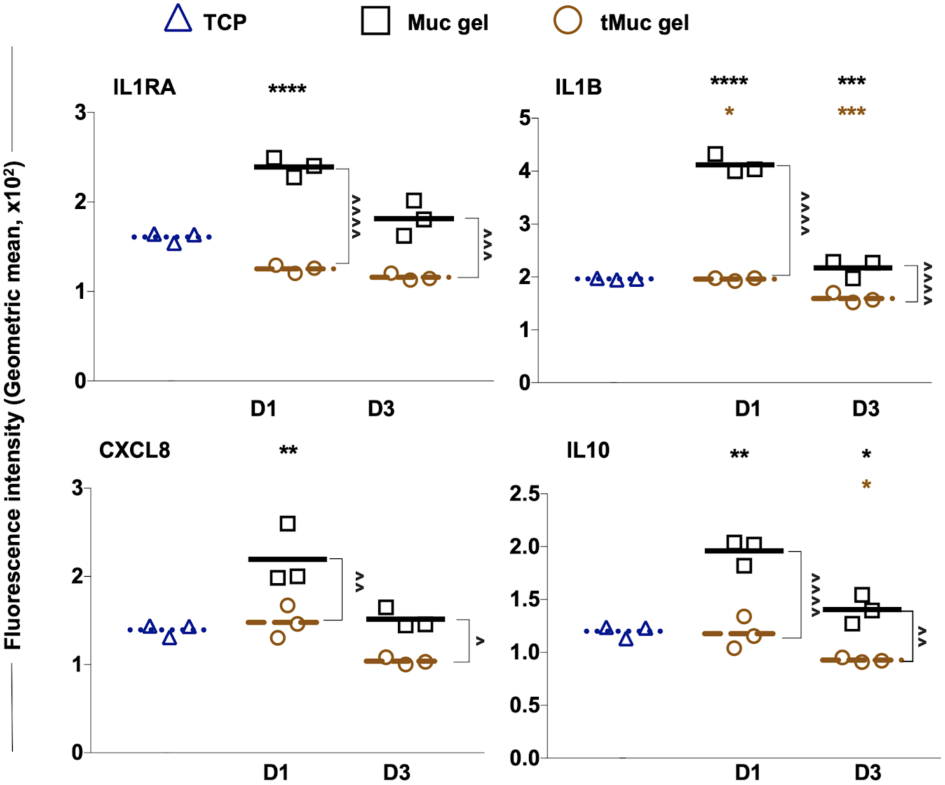
Intracellular cytokine expression at the protein level in THP-1 derived macrophages type 0 (THP1-M0) after being cultured on tissue culture polystyrene (TCP), Muc gels and Sialidase treated Muc gels (tMuc gel) on Day (D) 1 and 3, was analyzed by FACS. The data denote the mean of geometric mean of fluorescence intensity from three-independent experiments. Statistics was obtained by One-way ANOVA test by Prism 8.0. Black*, Brown*, and Black^ indicates the comparison between Muc gels vs TCP, tMuc gels vs TCP, and Muc gel vs tMuc gel individually. ‘*’, ‘**’, ‘***’, and ‘****’ indicate p values of < 0.05, 0.01, 0.0005, and 0.0001, respectively.

### Contact with Muc gels decreased the phagocytic ability of THP1-M0 while maintaining their endocytotic ability unchanged

In addition to major changes in the expression of cell markers and cytokines, the polarisation of macrophages also results in functional differences. In particular, the tendency of macrophages to uptake foreign objects by either endocytosis or phagocytosis has been associated with their polarization of macrophages *in vitro*^*27*^. We thus investigate the phagocytosis and endocytosis capacities of THP1-M0 after culturing them on TCP, Muc gel, or tMuc gel for 1 day. We show the Muc gel dampened the phagocytic activity of M0 but did not change their endocytic activity (**Figure 7**). Cells cultured on tMuc gel showed a similar trend but with a less pronounced decrease in the phagocytic activity.

**Figure 7.**
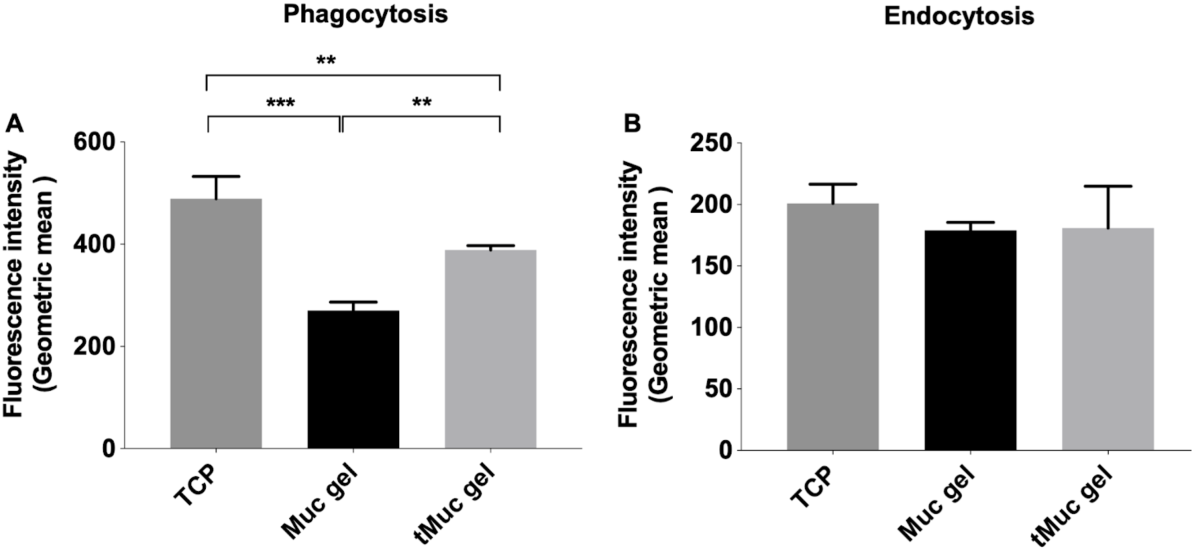
Phagocytosis and endocytosis of THP1-M0 cells cultured on tissue culture polystyrene (TCP), Muc gels and Sialidase treated Muc gels (tMuc gel) for one day and analyzed by FACS. Cells were treated with pHrodo green *E*. coli bioparticles to measure phagocytosis (A) and fluorescein labeled dextran (10 kD, sigma-aldrich) to measure endocytosis (B). Data reflect three independent experiments. Statistics were obtained via one-way ANOVA test among cells cultured on three different surfaces. ‘*’, ‘**’, ‘***’, and ‘****’ indicate p values of < 0.05, 0.01, 0.0005, and 0.0001, respectively.

### Macrophages type 0 derived from human peripheral blood monocyte cells (hPBMC-M0) are also activated by Muc gels in a sialic acid dependent manner

Although the protocol used to obtain macrophages from THP1 monocytes has been optimized to generate macrophages best resembling primary monocyte-derived macrophages, there persist differences in how they respond to stimuli^38^. We thus studied the response of human peripheral blood monocytes derived macrophages type 0 (hPBMS-M0) when cultured on Muc gels to increase the further validate the biological relevance of the results presented above. We sorted human monocytes (CD3^-^CD19^-^CD14^+^) by FACS based on cell surface markers (**Figure SI7**). The monocyte-macrophage differentiation was performed by incubation with macrophage colony-stimulating factor (M-CSF). The differentiated macrophages became adherent and expressed macrophage markers (**Figure SI8**). hPBMC-M0 cells were cultured on three different surfaces TCP, Muc gels, and tMuc gels over a period of 7 days. hPBMC-M0 were elongated on TCP on day 1, 3, and 7, spindle-shaped on Muc gels, and round with dendrites on tMuc gels (**Figure 8**). The cell-cultured on Muc gels and tMuc gels could be detached by pipetting, indicating a rather weak adhesion.

**Figure 8.**
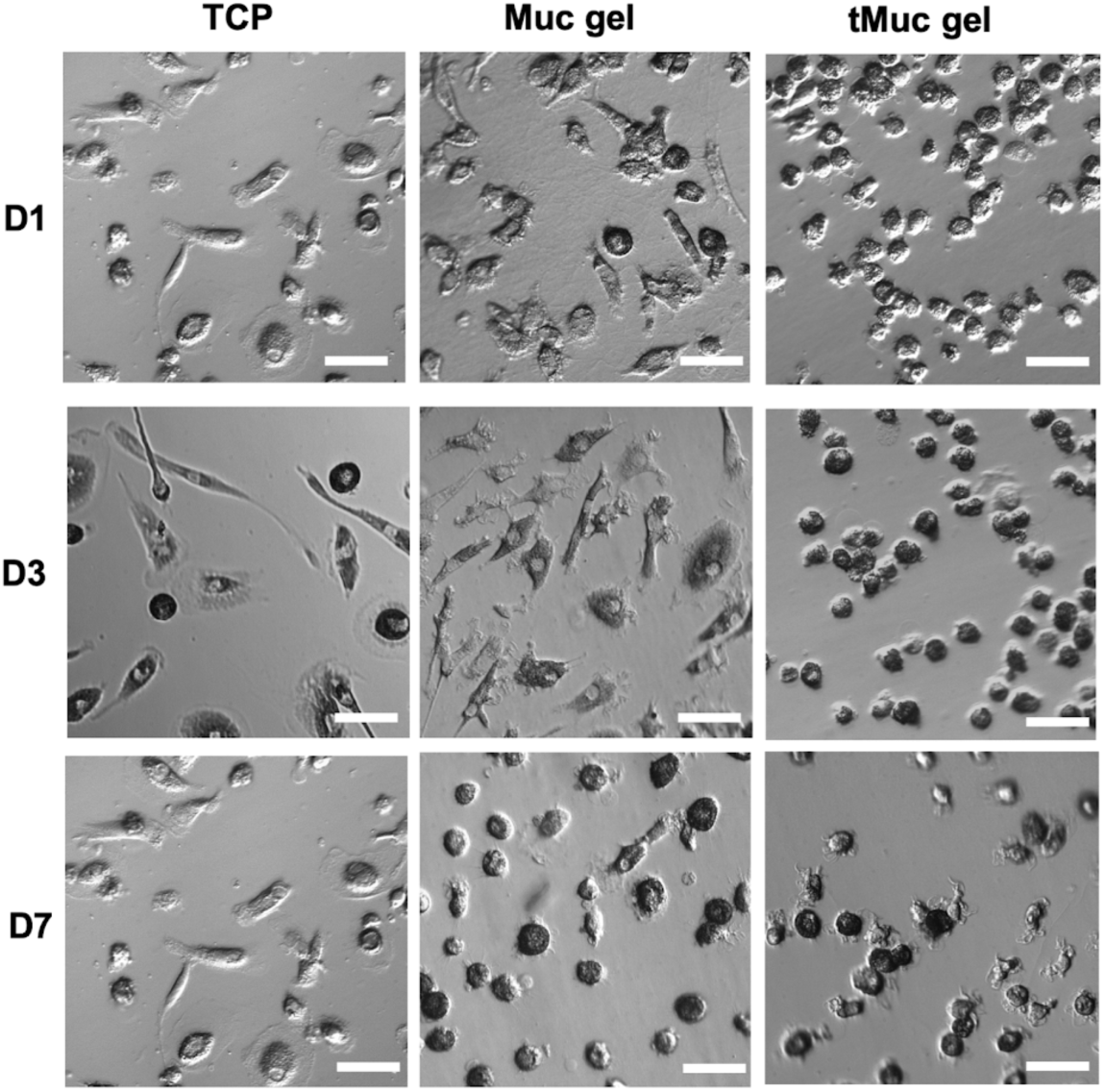
Representative phase-contrast images of hPBMC-M0 cultured on tissue culture polystyrene (TCP), Muc gel, and tMuc gels on day (D) 1, 3, and 7. Scale bar = 50 µm.

We investigated the expression of seven cytokines in hPBMC-M0 cells on day 1, 3, and 7. We show that both pro-inflammatory cytokines (*CXCL8, TNFa, CCL2, IL1B, VEGFA*) and anti-inflammatory (*IL1Ra, IL-10*) cytokines were significantly upregulated in cells cultured on Muc gels on day 1, then downregulated on day 3 and 7 (**Figure 9**). The partial removal of sialic acids in tMuc gels dampened the transient upregulation of all cytokines down to the levels in cells cultured on TCP. However, for *CXCL8* and *IL1B* the effect was limited, and expression levels were found to be significantly different from cells cultured on TCP. The results were similar for hPBMC-M0 cells derived from two independent donors and were consistent with the gene expression pattern of THP1-M0 cells.

**Figure 9.**
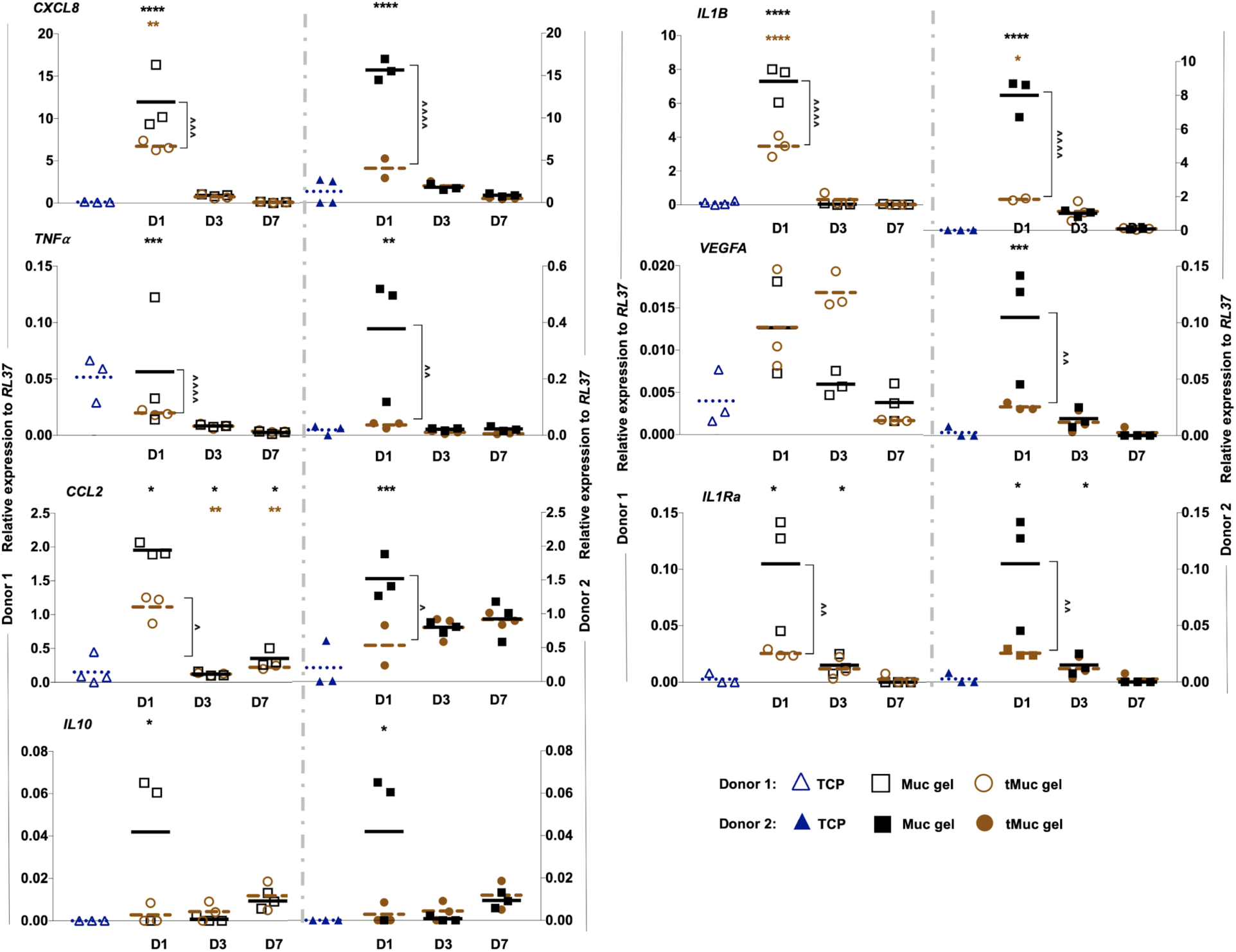
Gene expression in human peripheral blood monocytes-derived macrophages type 0 (hPBMC-M0) after being cultured on tissue culture polystyrene (TCP), Muc gels and Sialidase treated Muc gels (tMuc gel) on Day (D) 1, 3 and 7. The data denote the mean of relative gene expression to RPL-37 obtained from three independent repeats with duplicates. Donor 1, 2 are hollow and filled shapes respectively. Statistics was obtained by One-way ANOVA test by Prism 8.0. Black*, Brown*, and Black^ indicates the comparison between Muc gels vs TCP, tMuc gels vs TCP, and Muc gel vs tMuc gel individually. ‘*’, ‘**’, ‘***’, and ‘****’ indicate p values of < 0.05, 0.01, 0.0005, and 0.0001, respectively.

## Discussion

To investigate the relative short-term immune-modulatory effect of Muc gels, we cultured both cell line and primary cell-derived macrophages for 7 days at the surface of the hydrogels. *In vitro* study allows us to investigate the role of mucin glycosylation in the immunomodulatory effect in controlled conditions. We were able to modulate the sialic acid content of mucins while maintaining their ability to form crosslinked hydrogels. The resulting hydrogels have similar bulk elastic moduli (**Figure 2**) and nanomechanical surface properties (**Figure 3**) regardless of the sialic acid content, suggesting that sialic acid residues on mucins are not crucial for the formation of the crosslinking knots of Muc gels. We hypothesize that this is because the EDC/NHS chemistry applied to graft the Tz and Nb functionalities mainly targets carboxylic groups on the mucin-protein backbone^26^. The localization of Tz/Nb functionalities on the protein backbone is also supported by the absence of Tz and Nb ^1^H-NMR peaks in the glycan fraction after their removal from mucins by beta-elimination, while strong Tz and Nb ^1^H-NMR peaks were detected from the protein fraction (**Figure SI1**). However, since beta-elimination does not remove all mucin glycans and sialic acid residues, it is still possible that the absence of Tz and Nb in the glycan fraction is explained by their exclusive localization on glycans resistant to beta-elimination.

The similarities between Muc and tMuc gels in their mechanical properties and the levels of endotoxin and DNA impurities (**Figure SI9**) ensure that the effects observed on macrophages are solely due to the removal of sialic acid. Indeed the mechanical properties of the cellular substrate can affect a host of cell processes^39^ and impurities in the material alone can activate macrophages^40,41^. Such glyco-modulation of mucins with glycolytic enzymes prior to the material formulation could also be expanded to other enzymes, including glycosyltransferases. This, in turn, could enable the generation of a library of mucin materials that can be used to investigate the role of specific glycosylation patterns on the bioactivity of contacting bacterial or mammalian cells.

Using the Muc and tMuc gels, we characterized the interactions of non-polarized macrophages, both THP1-M0 and hPBMC-M0, with the materials’ surface. The Muc gels have already shown to be non-cytotoxic and support the survival of cells seeded on or in the hydrogels for several days^26^. Both THP1-M0 and hPBMC-M0 poorly adhered to the surface of Muc and tMuc gels. And although hPBMC-M0 spread and did not form clusters, pipetting alone was sufficient to detach them from the surface. This did not come as a surprise, as the materials carry similarities with other poorly cell-adherent materials such as alginate^42^ and hyaluronic acid^43^ that are also hydrated, negatively charged and do not carry any known binding ligands to integrins. This is also in agreement with previous reports of mucin coatings preventing cell adhesion^44,45^.

Then, with the objective to characterize the possible polarization resulting from contact with mucin materials, we measured cell surface markers and inflammatory cytokine expression levels. The primary hallmark is a general activation of cytokine gene expression and protein production when in contact with the Muc gels, followed by a reduction on day 3 and back to baseline on day 7. Both THP1-M0 that cluster rapidly into multicellular structures and the hPBMC-M0 that stay in direct contact with the substrates throughout the 7 days showed similar transient activation, suggesting that the activating signals were not strongly dependent on a constant contact with the material. It could be that the cells received an initial cue from the material, followed by material-independent regulation of the activation.

Beyond the expression of markers, THP1-M0 on Muc gels changes their function of phagocytosis significantly (**Figure 7A**) but not their endocytic capacity (**Figure 7B**). Interestingly, M1 macrophages polarized by IFNγ or LPS and IFNγ were shown to decrease their phagocytic ability compared to M0^27^, which is consistent with our results. However, there is not sufficient evidence to conclude that contact between M0 macrophages and a Muc gel would result in an M1 phenotype. Thus, neither the expression of cytokines, cell marker or the functional signature of macrophages suggests that M0 macrophages polarize toward M1 or M2 after contact with Muc or tMuc gels. The activation of macrophages described herein and in previous work^26^ support the existence of a continuum of macrophage phenotypes between M1 and M2^46^, especially *in vivo*^*47–49*^. Indeed, the details in how macrophages are activated depend on numerous factors such as the combination of biochemical signals, cell type, and kinetics. Complex environments encountered *in vivo*, or contact with complex materials such as Muc gels, would impact several of these factors.

A key result from this study is that sialic acid residues are important for the immune-modulating properties of mucin hydrogels. The bovine submaxillary mucins used herein contain a substantial amount of sialic acid residues (∼30%)^50^, which are mostly attach to N-acetyl-galactosamine via an alpha 2-6 bond (Sialyl-Tn antigen). Sialic acid is estimated to be one of the most abundant sugar in the mammalian glycome, with an estimated 8 % of all sugars being sialic acid, and their prevalence being up to 26 % amongst terminal sugars^51^. These highly accessible sugars bearing negative charges contribute to the physical properties and biological activities of mucins^52^. In immunity, sialic acids physically mediate cell-cell interactions and mask antigens, and biologically regulate the activation of the complement, leukocyte trafficking, and the activity of B cells through binding to sialic acid-binding receptors (*i.e.*, lectin proteins) on the immune cells^53^. In dendritic cells, neutrophils and macrophages, sialic acid–binding immunoglobulin-like lectins (siglec) regulate cytokine expression by inhibiting the toll-like receptor signaling pathway when bound to sialic acid residues of mucins^54,55^ and other ligands^56^. Although sialic acid was not completely absent in tMuc gels, a relative decrease compared to Muc gel still led to drastic changes in the macrophages’ response to the material. This could be an effect of lower ligand density and accessibility after sialidase treatment, which could hinder the binding of sialic acid receptors on macrophages that regulate their activities. Although sialic acid seems to be a critical component of Muc gel immunomodulation, this does not exclude the role of other sugars. Indeed, tMuc gels with low sialic acid contents activated *IL10* and down-regulated *MCR1, CCL2* (**Figure 5** and **9**, brown*) compared to unmodified Muc gels.

## Conclusion

In this study, we characterized the short-term response of macrophages to Muc gels and investigated the role of sialic acid in the bioactivity of the material. We were able to modulate the glyco-composition of mucin hydrogels without altering their bulk rheological properties and nanomechanical surface properties. We show Muc gels transiently activate macrophages in a sialic acid-dependent manner. Macrophages exposed to Muc gels could not be classified as M1 or M2, but showed broad expression of cytokines on day 1 followed by a decrease on day 3 and 7, with only a few exceptions. How these macrophage activation patterns translate into the broader immune reaction to implantation is unclear. In part because macrophages expression patterns and biomaterial implant outcomes are not well correlated and in another part because of the absence of many other immune components in our *in vitro* system. However, the low cytokine expression could be correlated with the low cytokine expression levels found 14 and 21 days after implantation of Muc gels in the intraperitoneal space of mice, which could be linked with high expression levels of cytokine inhibitor proteins^26^. This study also demonstrates that the glyco-modulation of crosslinkable mucin building blocks serves as a valuable tool to study the bioactivities of mucin materials. Such an approach could be expanded to establish a series of mucin hydrogel variants to study the interplay between glycan composition and cell response. For instance, this study highlights the importance of sialic acid in immune-modulating properties of Muc gels and suggests sialic acid immobilized on a backbone polymer could be a good candidate for artificial mucins recapitulating some of their intrinsic immune-modulating properties^57^.

## Materials and methods

### Materials

Tetrazine amine (Tz) and norbornene amine (Nb) were purchased from Bioconjugate Technology Company and TCI EUROPE N.V., repetitively. All chemicals were obtained from Sigma Aldrich. Cell culture medium and PCR related reagents were purchased from ThermoFisher Scientific. RNA extraction micro or mini kits were purchased from Qiagen. Human monocytes (THP-1) were purchased from ATCC, and human peripheral blood was purchased from a blood bank at the Karolinska university hospital.

### Synthesis of mucin Tz and Nb derivatives

We introduced Tz and Nb crosslinking functionalities onto mucins (Muc-Tz, Muc-Nb) as described before^26^. In brief, mucin was pre-dissolved in MES buffer (0.1 M MES, 0.3 M NaCl, and pH 6.5) at a concentration of 10 mg/mL. 1-ethyl-3-(3-dimethylaminopropyl) carbodiimide (EDC; 4 mmol per gram of dry mucin) and N-hydroxysuccinimide (NHS; 4 mmol per gram of dry mucin) were then added, and stirred for 15 min at room temperature. To the mixture, tetrazine (1 mmol, per gram of mucin) and norbornene (2 mmol, per gram of mucin) were added individually. The reaction mixtures were stirred at 4 °C overnight. After reaction, the reaction mixtures were dialyzed in 100 kDa cutoff-dialysis tubing for 2 days against 300 mM NaCl followed by dialysis against MilliQ H_2_O for 1 day. Samples were freeze-dried, and stored in −20 °C. Specifically, samples used for cell culture were filtered by a syringe filter (0.45 µm), then transferred into tissue culture flat tubes (screw cap with filter, 0.2 µm) for lyophilization to keep them sterile.

### Glycan modification and characterization

The sialic acid removal assay was conducted by using Neuraminidase immobilized on slurry (GlycoCleave Neuraminidase kit, GALAB technologies). Briefly, the gelling components of Muc-Tz and Muc-Nb were dissolved separately in a sodium acetate buffer (Sodium acetate 0.05mM, CaCl_2_ 1 mM, pH 5.5) at a concentration of 25 mg/mL. The solution was then mixed with 1mL Neuraminidase slurry and incubated overnight at 37°C at 30 rpm. To separate the Neuraminidase slurry and the enzyme-treated mucin derivatives, the mixture was passed through a 10 µm filter. The slurry was washed twice with an acetate buffer. After that, the flow-through was loaded into an Amicon Ultra-30K filter and then centrifuged at 4,000 g for 30 min to separate the enzyme-treated mucin and other small molecules. Next, 15 mL MQ H_2_O was added to the mucin fraction and then centrifuged at 4,000 g, 30 min three times to desalt the solution. Sterilization was performed by a syringe filter (0.45 µm), then the samples were loaded into tissue culture tubes equipped with screw cap with filter (0.2 µm). Samples were freeze-dried and stored at −20°C. The removal efficiency of sialic acid was investigated by anion exchange chromatography-based assays. In brief, Neuraminidase treated mucins were further treated by sulfuric acid to cleave all the glycans. Non-treated mucins were also treated by sulfuric acid and were used for quantification of the sialic acid content of mucin.

Sialic acid quantification was conducted by using the anion exchange chromatography-based assay as described above.

### Rheological characterization of Muc gels

Rheological measurements were performed using a research-grade shear rheometer (MCR302, Anton Paar) equipped with a plate-plate measuring geometry (measuring head: PP25, Anton Paar, Graz, Austria). The gap between the measuring head and the bottom plate (P-PTD200/Air, Anton Paar) was set to *d* = 150 µm for all measurements. Immediately before a measurement, the two components (Muc-Tz and Muc-Nb) of either the Muc gel or the tMuc gel were diluted in the particular buffer to a concentration of 25 mg/mL each. As a buffer solution, either PBS (pH = 7.4) or a mixture of RPMI 1640 (R8758, Sigma Aldrich, St Louis, Missouri, USA) containing 10 % FBS (F9665, Sigma Aldrich) and 1 % Penicillin and Streptomycin (P4333, Sigma Aldrich) was used. The two components were thoroughly mixed and centrifuged to remove bubbles before 100 µL of the sample were pipetted onto the rheometer plate. First, gel formation was analyzed for a total time span of *t* = 100 min. Both, the storage (*G*’) and loss modulus (*G*’’) were determined by a torque controlled (*M* = 5 µNm) oscillatory (*f* = 1 Hz) measurement. Afterwards, a strain-controlled frequency sweep (from *f*_start_ = 10 Hz to *f*_end_ = 0.01 Hz) was performed to determine the frequency dependent viscoelasticity of the cross-linked sample. For this frequency sweep, a constant strain was used, which was chosen as the average of the five last values determined in the torque-controlled gelation measurement.

### Surface nanomechanical properties of gels by atomic force microscopy

Silicon wafers (22 × 22 mm) were cleaned using a 2 % Deconex solution (Borer Chemie AG, Switzerland) in a sonicator for 15 min, then rinsed with MilliQ water, and ethanol sequentially, and dried using a filtered nitrogen jet. The gelling components for Muc gels and tMuc gels were premixed and deposited on substrates separately and incubated for 1 h in a humidified chamber to allow proper gelation. The nanomechanical measurements were conducted in force volume mapping mode using a JPK NanoWizard 3 atomic force microscope (JPK Instruments AG, Berlin, Germany). Before the measurements, a drop of PBS is loaded onto the gel to have an aqueous phase. The EBD biosphere B100-CONT (Nanotools) probe of a well-calibrated and measured sphere tip outer radius of 100 nm and a measured spring constant of 0.26 N/m was used for nanoindentation measurements. The acquired force curves were analyzed using standard JPK data processing software (JPK, version 6.1.86). The Derjaguin-Muller-Toporov (DMT) model was fitted to determine the elastic modulus on approaching and retracting force curves following a previous publication^33^. The force volume maps were measured on an area of 2×2 µm^2^ with 8 by 8 data points. The applied normal force was 0.3 nN, and the acquisition speed was 4 µm/s. Three different areas of 2×2 µm^2^ were measured in order to evaluate the average nanomechanical parameters. We observed differences in the nanomechanical surface properties between the approach and retract mapping regimes for both Muc gels (**Figure 3A, B**) and tMuc gels (**Figure 3A, B**). These are possibly due to the different contributions in each regime, with a combined elastic and viscous contributions in the approach maps, and a predominant elastic contribution in the retraction maps. This also indicates the importance of analysing both approaching and retraction regimes in AFM force volume mapping for soft materials, the surface dynamics of which should be taken into account and a complicated tip-surface interaction occurs in the measurements^33,58^. Moreover, commonly applied contact mechanics models such as Hertz and/or the Derjaguin-Muller-Toporov (DMT) are limited for studying soft gels due to the substantial viscous contribution from those soft gels. We thus also evaluated surface stiffness parameters, which do not require any contact mechanics model fitting and can be more suitable for the direct comparison of the nanomechanical surface property of Muc gels and tMuc gels using the same AFM probe.

### THP-1 cell cultivation and differentiation

Human monocytes THP-1 were purchased from ATCC and cultured in RPMI-1640 medium supplemented with 10% FBS, and penicillin/streptomycin (100 U/mL). Cells were split at the ratio of ⅕ when the cell density reached 1 × 10^6^ cells/mL. To differentiate cells into macrophage type 0 (M0), the THP-1 cells were cultured in the culture medium used above and supplemented with 150 nM phorbol 12-myristate 13-acetate (PMA, Sigma-Aldrich) for 72 hours, followed by 24 h incubation in a complete cell culture medium without PMA. To confirm the differentiation of THP-1, the cell morphological change was examined under bright field microscope and macrophage markers CD36 (2.5 µg per 1×10^6^ cells in 100 µL; Cat. no. 108418, BioLegend) and CD71 (2.5 µg per 1×10^6^ cells in 100 µL; Cat. no. 108418, BioLegend) were evaluated by FACS.

### Human monocytes isolated from peripheral blood and differentiation

Human monocytes were isolated from human peripheral blood from 2 donors purchased from Blood Bank at Karolinska Sjukhuset. Mononuclear cells were acquired by using Ficoll-Paque PREMIUM density gradient media (GE healthcare Life Science) according to the instruction. Briefly, blood was diluted with PBS at ratio of ⅘, which then was carefully layered onto the Ficoll-Pague media at the ratio of ⅘. To obtain the mononuclear cell, the samples were then centrifuged at 700 g for 40 min with acceleration and deceleration speed level at 4. The serum was sterilized by using 0.45 µm filters and stored at 4°C for further usage. The mononuclear cells were washed in PBS and centrifuged for 10 min at 700g to remove the Ficoll media. The cells were cleaned through a 70 µm cell strainer (Corning) to get rid of clumps and then counted using a bürker chamber. Monocytes were enriched by using monocytes enrichment kit (BD biosciences) according to the manufacture instructions. The cells were resuspended in an IMAG™ buffer solution and incubated with the monocyte enrichment cocktail and CD41 antibodies at a concentration of 5 µL per 1×10^6^ cells for 15 min. The non-conjugated antibodies were washed away by IMAG™ buffer, the cell pellet was then resuspended in IMag ™ streptavidin Particles Plus-DM at the concentration of 5 µL per 1×10^6^ cells for 15 min. The enriched monocytes fraction was negatively selected and further sorted by FACS. Briefly, the monocytes were incubated for 10 min at room temperature with human BD Fc-block (2.5 µg per 1×10^6^ cells in 100 µL, Cat. no. 564220, BD biosciences). The cells were further incubated with the following antibody cocktail for 30 min at 4°C: APC-H7 Mouse Anti-Human CD3 Clone M-A712 (2.5 µg per 1×10^6^ cells in 100 µL, Biolegends), PE Mouse Anti-Human CD14 Clone M5E2 (2.5 µg per 1×10^6^ cells in 100 µL, Biolegends) and BB515 Mouse Anti-Human CD19 Clone HIB19 (2.5 µg per 1×10^6^ cells in 100 µL, Biolegends). Cells were washed with 5 mL PBS and then resuspended in 5mL PBS containing 20% serum. The cells within the gate of CD3-CD19-CD14+ were then sorted using FACS.

To differentiate the monocytes into M0, monocytes were cultured in RPMI-1640 medium supplemented with 20% of endogenous serum, penicillin/streptomycin (100 U/mL), and macrophage colony-stimulating factor (M-CSF, Gibco, 1µg per 5mL medium, Cat. no. PHC9501, Gibco^**®**^) in a T-25 culture flask for 5 days.

### Gene expression analysis by real-time PCR

Total RNA of cells was extracted by using either Qiagen RNeasy mini kit or Qiagen RNeasy micro kit depending on cell numbers obtained. The extracted mRNA was diluted to a concentration of 0.67ng/µL and synthesized into cDNA using Superscript III polymerase (Invitrogen). Real-time PCR was then performed to analyze the gene expression by using a TaqMan Gene Expression Master Mix (Thermo Fisher Scientific) together with TaqMan probes. See the Taqman probes in the supporting information **Table S1**. The RT-PCR were carried out in a CFX96 Touch™ Real-Time PCR Detection System (Bio-Rad) with the following cycling conditions: 50°C for 2 min, 95°C for 10 min, 95°C for 15 sec, 60°C for 1 min, and then go to step 3 for 50 cycles. RPL37 was used for Thp1 derived macrophages, while ACTB were used as housekeeping gene for primary monocytes derived macrophages and cells from the *in vivo* study.

### Intracellular cytokine expression by FACS

THP1-M0 cells cultured on TCP, Muc gels, and tMuc gels were incubated with brefeldin A buffer (diluted to 1x with complete cell culture medium, Cat. no. 420601, Biolegend) for 5 hours. Cells were then harvested, washed with a washing buffer (PBS containing 0.5% bovine serum albumin (BSA) and 0.1% sodium azide) twice, and then resuspended in a FACS™permeabilizing solution (Cat. no. 347692, BD bioscience) for 10 min at room temperature. After permeabilization, cells were washed with 1 mL of a washing buffer and centrifuged at 500 g for 5 min. Cells pellets were then incubated with 500 µl of 1% paraformaldehyde at room temperature and then washed twice with a washing buffer. Cells were then incubated with antibody cocktail for 30 min on ice, containing anti-IL1RN (10 µl per 1×10^6^ cells in 100 µL, Cat. no. 340525, BD bioscience), anti-IL1B (5 µl per 1×10^6^ cells in 100 µL, Cat. no. 340515, BD bioscience), anti-IL10 (5 µl per 1×10^6^ cells in 100 µL, Cat. no. 562400, BD bioscience), and anti-IL8 (5 µl per 1×10^6^ cells in 100 µL, Cat. no. 563310, BD bioscience). Cells were then washed, resuspended in the washing buffer before being subjected to FACS analysis.

### Phagocytosis and endocytosis

pHrodo green *E*. coli bioparticles (LifeTech) and Fluorescein labeled dextran (10 kD, sigma-aldrich) were used to investigate the phagocytosis and endocytosis function of THP-1 derived M0. Briefly, Cells were seeded on TCP, Muc gels, and tMuc gels. After one day, cells were then incubated with either dextran or E. coli bioparticles (5 µg/ml) for 60 min at 37°C. Cells without treatment served as negative control. The internalization of the particles was then quantitatively measured by the geometric mean of fluorescence intensities (GMFI) using flow cytometry.

## Statistical analysis

Data are shown as mean of three independent experiments. The significance was analyzed via non-parametric one-way ANOVA test using GraphPad Prism 8.0; ‘*’, ‘**’, ‘***’, and ‘****’ indicate p values of < 0.05, 0.01, 0.0005, and 0.0001, respectively.

## Acknowledgement

T.C. financial support from the Swedish Foundation for Strategic Research (Grant No. FFL15-0072), FORMAS (Grant No. 2015-1316), and the Swedish Research Council (Grant No. 2014-6203). O.L. acknowledges financial support from the Deutsche Forschungsgemeinschaft (DFG) through project B11 in the framework of SFB863. The authors acknowledge Francisco Javier Vilaplana Domingo from KTH for assistance with sialic acid quantification.

