## Supplemental Figures and Tables for "Mucin hydrogel glyco-modulation to investigate immune activity of mucin glycans"

### Methods

#### $^1$H-NMR detecting Tz and Nb after beta-elimination

To identify whether Tz and Nb functionalities were on the mucin glycans or protein backbone, Beta-elimination kit (Sigma-Aldrich, GlycoProfile™ beta-Elimination Kit) was used to remove the O-glycans from Muc-Tz and Muc-Nb. The protein faction and glycan fractions were separated and the functionalities of Tz or Nb were detected by NMR. Muc-Tz and Muc-Nb were dissolved in MQ $H_2O$ at a concentration of 5 mg/mL individually. 200 µL of beta-elimination solution (per 1.5 mg Muc-Tz or Muc-Nb) was then added and incubated at 4 °C overnight.

To separate the glycans and mucin protein backbone, the mixture was loaded in an Amicon® Pro Purification System with 10 kDa Amicon® Ultra-0.5 Device and centrifuged at 4 000 g for 30 minutes at 4 °C. The flow through contains eliminated glycans was collected. To desalt the mucin protein backbone, Milli-Q water was added to the top solution in the filter tube followed by centrifugation at 4 000 g for 30 minutes at 4 °C. The top solution from the filter tube are the protein fraction, was collected. Both the glycans and protein backbone were lyophilized for further analysis.

Mucin derivatives were dissolved in deuterium oxide (Sigma-Aldrich). After dissolution, the 600 µL of the solutions were transferred into a 5 mm NMR tube (Norrel, USA). $^1$H-NMR spectra were obtained on a on a Bruker Ultrashield plus 500 MHz spectrometer (Bruker Corporation, USA). The processing for the spectra were carried out by using the MestReNova software (version 12.0.1-20560). The degrees of the functionalities were evaluated by comparing the ratio of integral Tz proton peaks (δ10.4, δ6.0, and δ5.2) or norbornene proton peaks (δ6.3-5.8) to protein backbone proton peak at δ0.92.

**Beta-elimination efficiency evaluation by anion exchange chromatography-based assay**

To know glycan removal efficiency on mucin by the ß-Elimination assay. After ß-Elimination, mucins were further treated with sulfuric acid (0.1 N $H_2SO_4$, 1 h, 80 °C) to de-*O*-acetylated and hydrolyze mucin, which was already shown effective to remove > 99% O-glycans[59]. After sulfuric acid treatment, the pH of the solutions was neutralized by the addition of NaOH (0.2 N). Neu5Ac (Sigma Aldrich) dissolved in MQ $H_2O$ was used as the standard and also subjected to the treatment as mucin samples. Samples were being loaded to a high performance anion exchange chromatography with pulsed amperometric detection (HPAEC-PAD) with a ICS-3000 system (Dionex) equipped with a CarboPac PA1 column (4 × 250 mm, Dionex)[60].

To calculate the sialic acid content, the integrations of sialic acid elution peaks from each sample and standards were obtained. The quantity of sialic acid of each sample was calculated based on a formula obtained from the standard curve of the sialic acid elution peak integration vs the concentration of sialic acid standards.

**Results**

In our previous study, we hypothesized Muc-gels crosslinking knots are primarily located on mucin protein backbone[26], due to the fact that the storage modulus of Muc-gels decreased gradually following the exposure to trypsin and unchanged sialic acid content among mucins, mucin-Tz, and mucin-Nb. In this study, we further attempted to locate the location of Tz and Nb functionalities by a combination of beta-elimination and $H^1$NMR assays. We first eliminated Mucin glycans from mucin-Tz and mucin-Nb, then analyzed the glycan and protein fractions by $H^1$NMR. We show $H^1$NMR peak for Tz and Nb are detected on mucin protein backbone fraction(**Figure SI1**), but not on the glycan fraction. We further treated protein fraction with sulphuric acid as indicated above to know the beta-elimination efficiency. We then analyzed sialic acid content. Unfortunately, we detected sialic acid elution peak in the anion exchange chromatography-based assay. This indicates that beta-elimination for removing mucin glycan is insufficient.

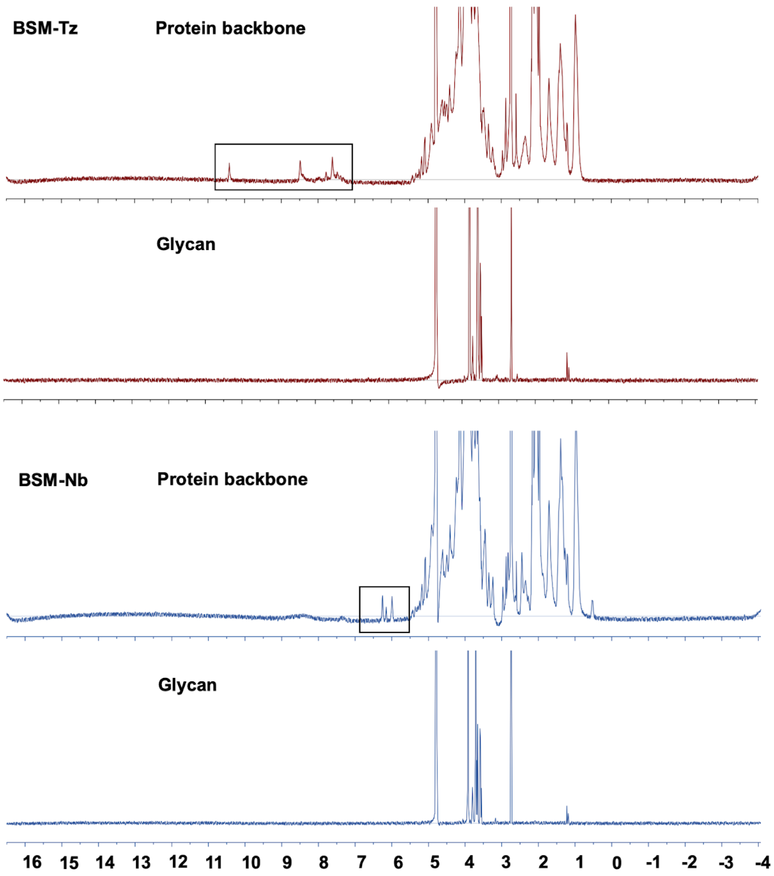


**Figure SI1.** $^1$H-NMR spectra of protein backbone and glycan fractions of mucins gelling components after the beta-elimination assay. Black boxes indicate the appearance of aromatic protons in Muc-Tz spectra and alkene protons in Muc-Nb individually in the protein backbone fraction. The spectra of non-functionalized mucins can be found in our previous publication[26].

We also solubilized Muc gel-geling components mucin-Tz and mucin-Nb in a complete cell culture medium, which are used to make the Muc gels and tMuc gels for cellular experiments in this study. We then studied the rheological behavior upon mixing with a time-sweep followed by a frequency sweep (**Figure SI2**). Our results show that both, the Muc-gel gelation time and the elastic modulus, were not impacted by the cell culture medium.

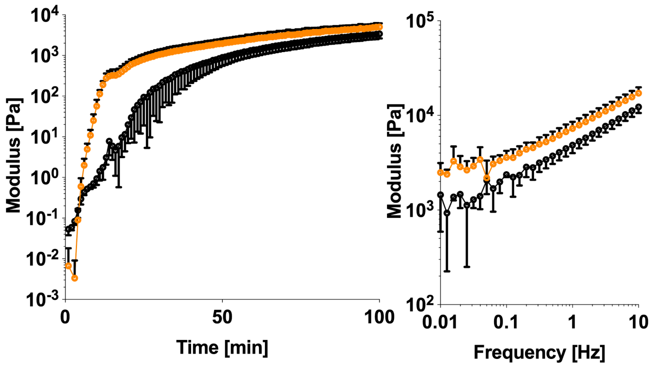


**Figure SI2.** Rheological characterization of Muc gels when using a cell complete medium as solvent for gelling components of Muc-Tz and Muc-Nb. Left: Time-dependent rheological measurements of the mixed Muc-Tz and Muc-Nb solution demonstrate the cell complete medium did not affect the crosslinking reaction. Right: Final frequency-dependent viscoelastic moduli of the crosslinked Muc-gel. The error bars denote the standard deviation as obtained from measurements of $n$ = 3 independent samples.

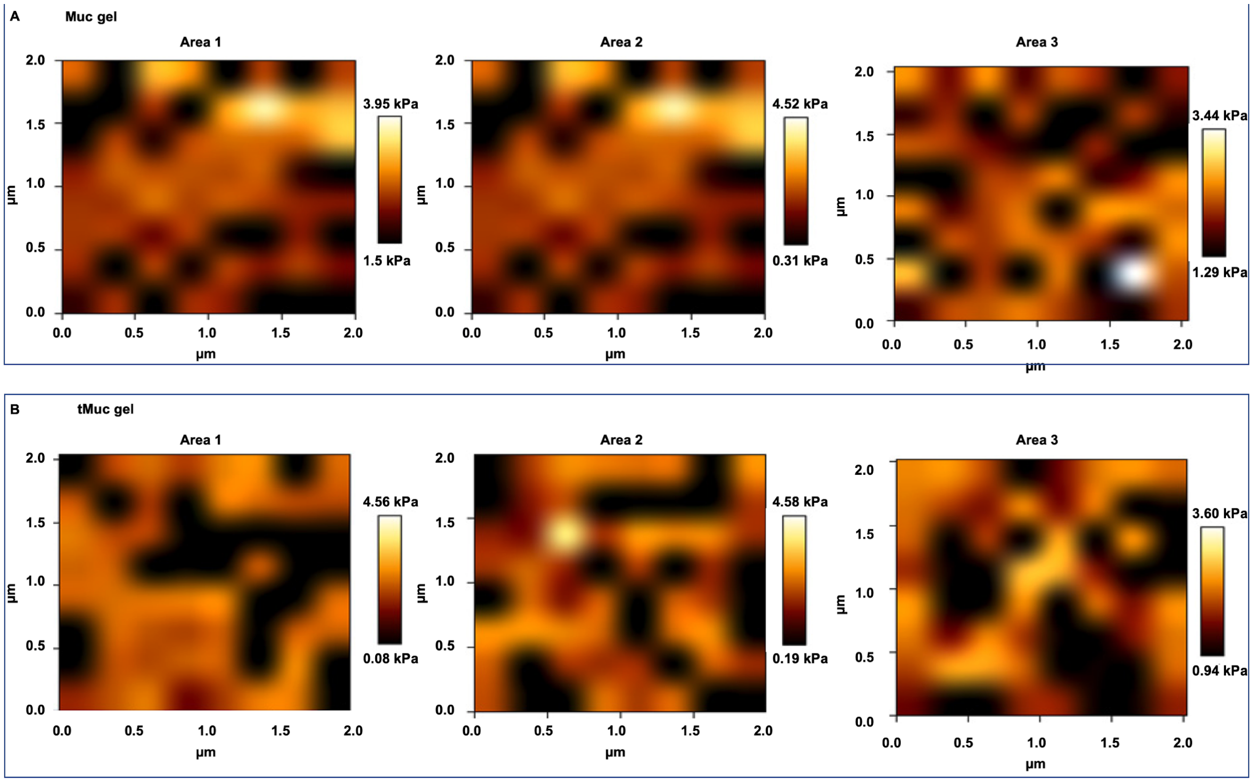


**Figure SI3** Approach mechanical mapping of the Muc gels (A) and tMuc gels (B) by AFM under force volume mapping mode.

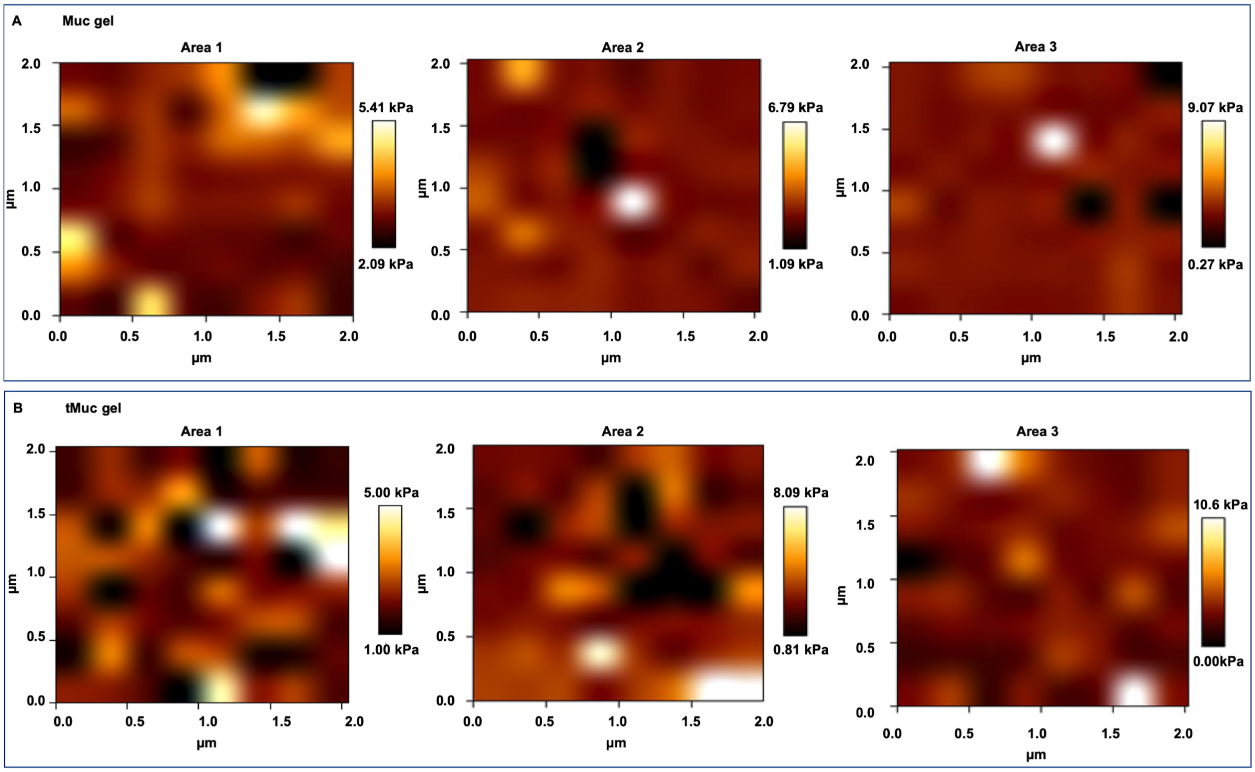


**Figure SI4** Retract mechanical mapping of the Muc gels (A) and tMuc gels (B) by AFM under force volume mapping mode.

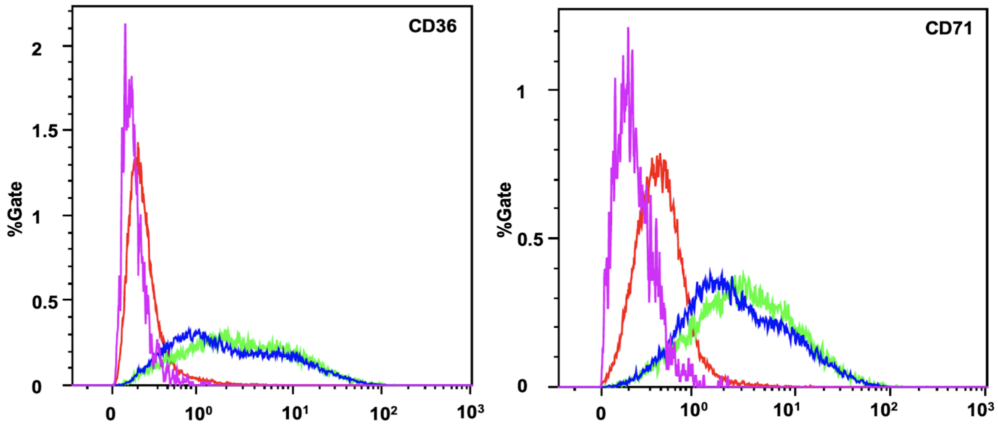


**Figure SI5.** Human monocytes THP1 derived macrophage type 0 (THP1-M0) upregulated macrophage markers (CD36 and CD71) when compared to THP1 as analyzed by FACS. Red and purple curves indicate THP1. Other curves indicate THP1-M0 from two independent experiments.

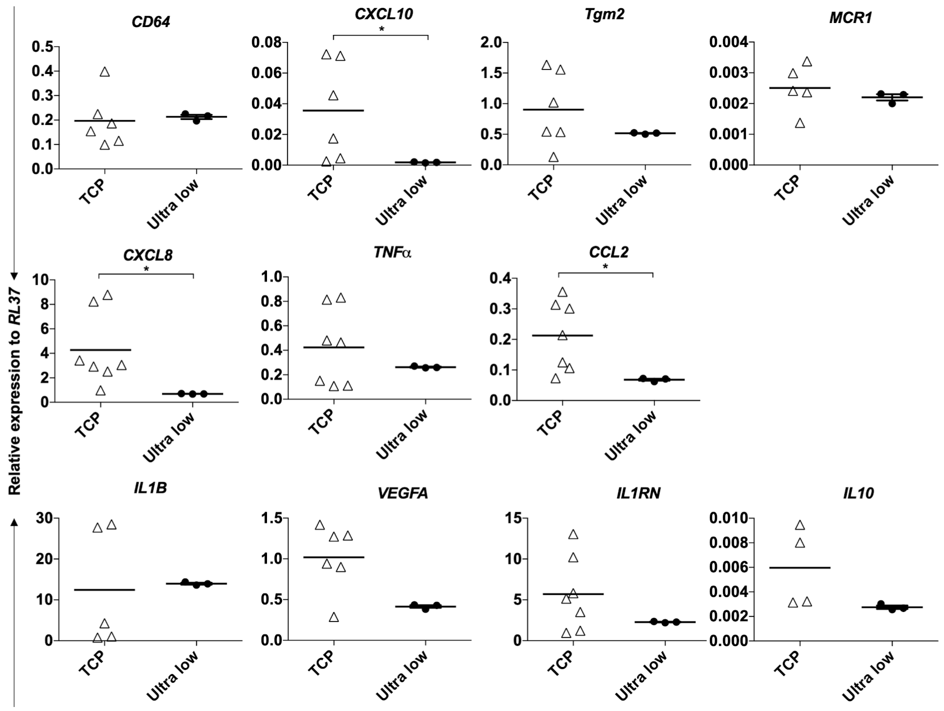


**Figure SI6.** Gene expression in THP1-M0 cultured on tissue culture polystyrene (TCP) and ultra-low surface (Ultra low) was analyzed by RT-PCR. The data denote the mean of relative gene expression to *RPL37* obtained from three independent repeats with duplicates. Statistics was obtained by One-way ANOVA test by Prism 8.0. '*', '**', '***', and '****' indicate p values of < 0.05, 0.01, 0.0005, and 0.0001, respectively.

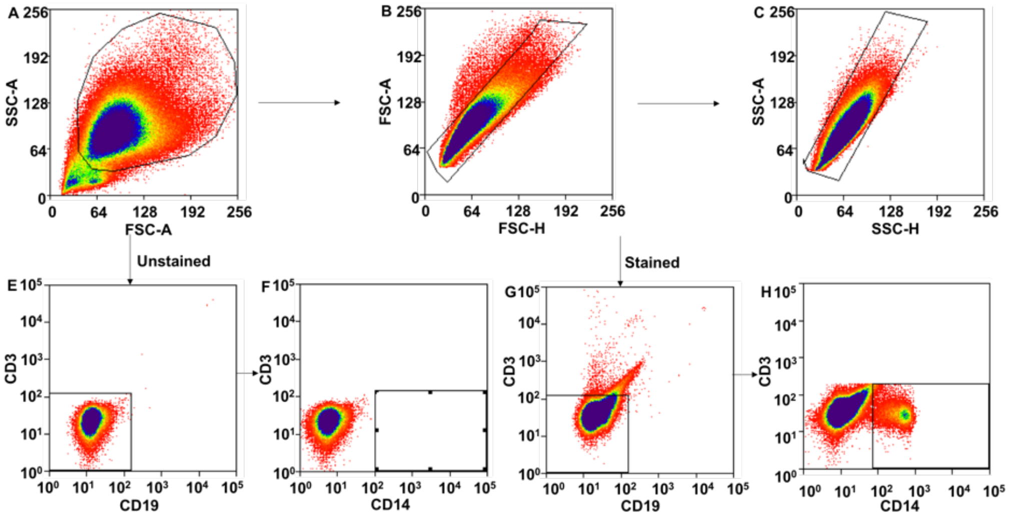


**Figure SI7**. Representative FACS profiles show the gating strategy for sorting monocytes ($CD19^-$ $CD3^-CD14^+$) from human peripheral blood mononuclear cell after monocytes negative enrichment. A-C are the cell populations. E and F are for unstained samples and G and H are for stained samples.

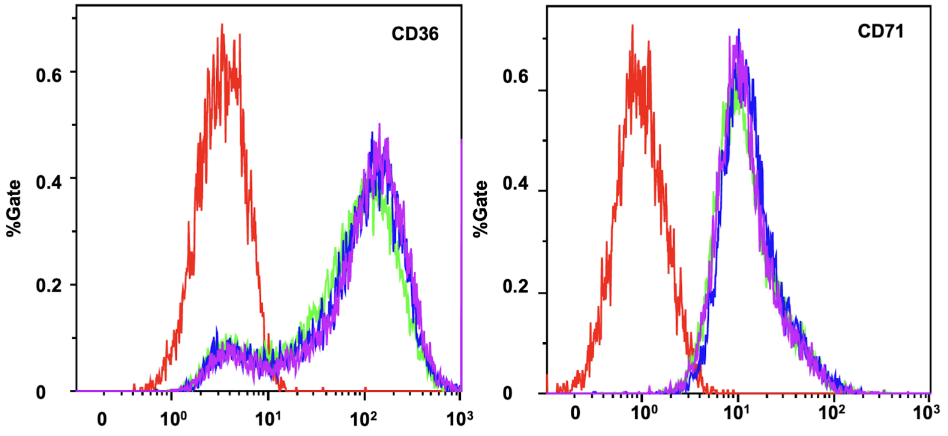


**Figure SI8**. Human peripheral blood monocyte cells derived macrophages type 0 (hPBMC-M0) upregulated macrophage markers (CD36 and CD71) when compared to monocytes as analyzed by FACS. Red curve indicates monocytes. Other curves indicate hPBMC-M0 from two donors.

Mucins preparation could be accompanied with contaminants such as LPS and DNA that could elicit some response from the macrophages. Both LPS and DNA content were quantified and showed that the treatment of neuraminidase did not change the LPS and DNA contents (**Figure SI2**). In addition the co-upregulation of both anti- and pro-inflammatory cytokines is not explained by the action of inflammatory contaminants alone[40], and the sialic-acid dependence of the effect point to the action of the mucins and its glycan.

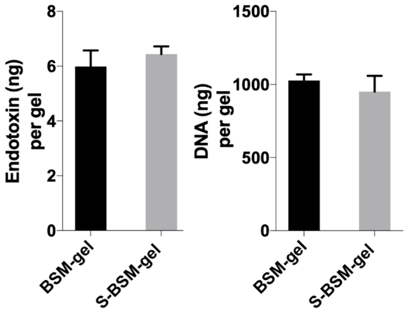


**Figure SI9.** Quantification of endotoxin and DNA contaminants in Muc gel and tMuc gel.

**Table SI1.** Taqman probes used for gene expression assays[61–63]

| Taqman probe | Assay ID | Classification |
|---|---|---|
| *CD64* | Hs00417598_m1 | M1 marker |
| *CXCL10* | Hs00171042_m1 | M1 marker |
| *Tgm2* | Hs01096681_m1 | M2 marker |
| *MCR1* | Hs00267207_m1 | M2 marker |
| *CXCL8* | Hs00174103_m1 | Pro-inflammatory and Anti-wound healing |
| *TNF-a* | Hs00174128_m1 | Pro-inflammatory and Anti-wound healing |
| *CCL2* | Hs00234140_m1 | Pro-inflammatory and Anti-wound healing |
| *IL1B* | Hs01555410_m1 | Pro-inflammatory and Pro-wound healing |
| *VEGFA* | Hs00900055_m1 | Pro-inflammatory and Pro-wound healing |
| *IL1Ra* | Hs00893626_m1 | Anti-inflammatory and Pro-wound healing |
| *IL10* | Hs00961622_m1 | Anti-inflammatory and Anti-wound healing |
